# A phasevarion controls multiple virulence traits, including expression of vaccine candidates, in *Streptococcus pneumoniae*

**DOI:** 10.1101/2022.02.08.479631

**Authors:** Zachary N. Phillips, Claudia Trappetti, Annelies Van Den Bergh, Gael Martin, Ainslie Calcutt, Victoria Ozberk, Patrice Guillon, Manisha Pandey, Mark von Itzstein, W. Edward Swords, James C. Paton, Michael P. Jennings, John M. Atack

## Abstract

*Streptococcus pneumoniae* is the most common cause of bacterial illness worldwide. Current vaccines based on the polysaccharide capsule (PCV-13 and PPSV-23) are only effective against a limited number of the >100 capsular serotypes. A universal vaccine based on conserved protein antigens requires a thorough understanding of gene expression in *S. pneumoniae.* Restriction-Modification (R-M) systems, classically described as a defence against bacteriophage, are almost ubiquitous in the bacterial domain, and roles other than phage defence. All *S. pneumoniae* strains encode the SpnIII R-M system. This system contains a phase-variable methyltransferase that randomly switches specificity, and controls expression of multiple genes; a phasevarion. We aimed to determine the role of the SpnIII phasevarion during pneumococcal pathobiology and determine if phase-variation resulted in differences in expression of protein antigens that are being investigated as vaccine candidates. Using ‘locked’ *S. pneumoniae* strains that express a single SpnIII methyltransferase specificity, we found significant differences in clinically relevant traits, including survival in blood, and adherence to and invasion of human cells. Crucially, we also observed differences in expression of numerous proteinaceous vaccine candidates, which complicates selection of protein antigens for inclusion in a universal protein-based pneumococcal vaccine. This study will inform future vaccine design against *S. pneumoniae* by ensuring only stably expressed candidates are included in a rationally designed vaccine.

**Significance Statement:** *S. pneumoniae* is the world’s foremost bacterial pathogen. *S. pneumoniae* encodes a randomly expressed epigenetic regulator, a phasevarion (phase-variable regulon), that results in random expression of multiple genes. Previous work demonstrated that the pneumococcal SpnIII phasevarion switches between six different expression states, generating six unique phenotypic variants in a pneumococcal population. Here, we show that this phasevarion generates multiple phenotypic differences relevant to pathobiology. Importantly, expression of conserved protein antigens varies with phasevarion switching. As capsule expression, a major pneumococcal virulence factor, is also controlled by the phasevarion, our work will inform the selection of the best candidates to include in a rationally designed, universal pneumococcal vaccine.

## Introduction

*Streptococcus pneumoniae,* the pneumococcus, is a human-adapted bacterial pathogen of global importance [1]. *S. pneumoniae* commonly colonises the nasopharynx asymptomatically in both children and adults [2]. Colonisation may develop into diseases of the respiratory tract, and lead to life-threatening meningitis and sepsis (invasive pneumococcal disease; IPD) [1]. *S. pneumoniae* is the predominant cause of middle ear infection (otitis media; OM), the most common bacterial infection in children (>350 million cases annually), and the leading cause of childhood antibiotic prescription and visits to healthcare providers in the developed world [3, 4]. *S. pneumoniae* is also one of the most common bacterial causes of exacerbations of chronic obstructive pulmonary disease (COPD). COPD affects >380 million people globally, costing $1 billion in healthcare annually [5]. In 2015 *S. pneumoniae* infection was responsible for approximately 1.5 million deaths from pneumonia alone, including 650 000 children aged <5 years [6]. The prevalence, morbidity, and mortality, associated with *S. pneumoniae* mean it is extremely important to understand the pathobiology of this organism to develop better treatments and vaccines. Current vaccines against *S. pneumoniae* are based on the pneumococcal capsule (conjugate vaccine PCV-13, and purified polysaccharide vaccine PPSV-23), only target a limited number of the >100 capsular serotypes [1, 7], and are only moderately effective at preventing OM [8]. A universal pneumococcal vaccine based on conserved protein antigens could potentially provide complete protection against all strains.

Restriction Modification (R-M) systems are almost ubiquitous in bacteria, where they were originally characterised as a defence against foreign DNA, typically from infection by bacteriophage [9]. In these systems, the restriction (R) enzyme cleaves foreign DNA at a particular DNA target sequence, whilst the cognate methyltransferase (modification; M) protects host DNA by methylating this same target sequence on host DNA [10]. In addition, many methyltransferases associated with R-M systems have a role in gene regulation. In a number of bacterial pathogens, the methyltransferase is phase-variable. Phase-variation is the random and reversible switching of gene expression, and is usually associated with bacterial surface features [11]. Phase-variable expression of DNA methyltransferases leads to different methylation patterns in a bacterial population, dependent on the phase-variable state of the methyltransferase in each individual bacterium in the population. These methylation differences influence expression of multiple genes epigenetically, in systems called phasevarions (<u>phase</u>-<u>vari</u>able regul<u>ons</u>) [12]. Phasevarions have been characterised in many bacterial pathogens such as *Neisseria spp* [13], *Moraxella catarrhalis* [14], *Haemophilus influenzae* [15, 16] and *Streptococcus suis* [17]. In every case, phasevarions regulate genes involved in pathobiology, and frequently genes encoding vaccine candidates. *S. pneumoniae* encodes a phase-variable Type I R-M system, the SpnD39III phasevarion [18], herein abbreviated to SpnIII. Type I R-M systems encode restriction *(hsdR),* modification *(hsdM)* and specificity *(hsdS)* subunits. The *hsdR* gene encodes the restriction endonuclease, while the *hsdM* gene encodes the cognate methyltransferase. The *hsdS* gene encodes a specificity subunit, HsdS, which determines the sequence cleaved or methylated by the HsdR and HsdM, respectively [10]. An R2M2S pentamer forms the active restriction enzyme, whilst an M_2_S trimer forms an active, stand-alone methyltransferase [10]. Each HsdS is composed of two target recognition domains (TRDs), with each TRD contributing half to the overall specificity of the complex [12]. A version of the SpnIII system is present in every publicly available genome of *S. pneumoniae* (>200 strains) [18]. The *hsdR* and *hsdM* genes of the SpnIII system are highly conserved between strains, and the methyltransferase of the SpnIII system is always expressed. The SpnIII locus encodes multiple, variable *hsdS* loci, and a recombinase. Homologous recombination between the variable *hsdS* loci (expressed *hsdS,* and downstream silent *hsdS’* and *hsdS”* genes) leads to multiple HsdS protein variants being expressed in a population. In strain D39 [18] and TIGR4 [19], this leads to six distinct DNA methyltransferase specificities being expressed in a pneumococcal population; SpnIII alleles A-F. These six different methyltransferase specificities result in six different gene expression profiles, resulting in six distinct phasevarions in a pneumococcal population (**Figure 1A**). Studies of the phenotypic effects of the SpnIII phasevarion in *S. pneumoniae* strains D39 and TIGR4 have found differential regulation of virulence determinants such as opacity, capsule and biofilm formation, commensurate with phase variation between methyltransferase specificities A-F [18, 19]. The presence of the SpnIII phasevarion complicates the identification of conserved antigens, as the full impact of this system on pneumococcal gene regulation and pathobiology has not been investigated. Understanding the influence of the SpnIII phasevarion on gross pneumococcal phenotype, and specific effects regarding expression of conserved protein antigens, is essential to designing successful treatments and vaccines.

**Figure 1.**
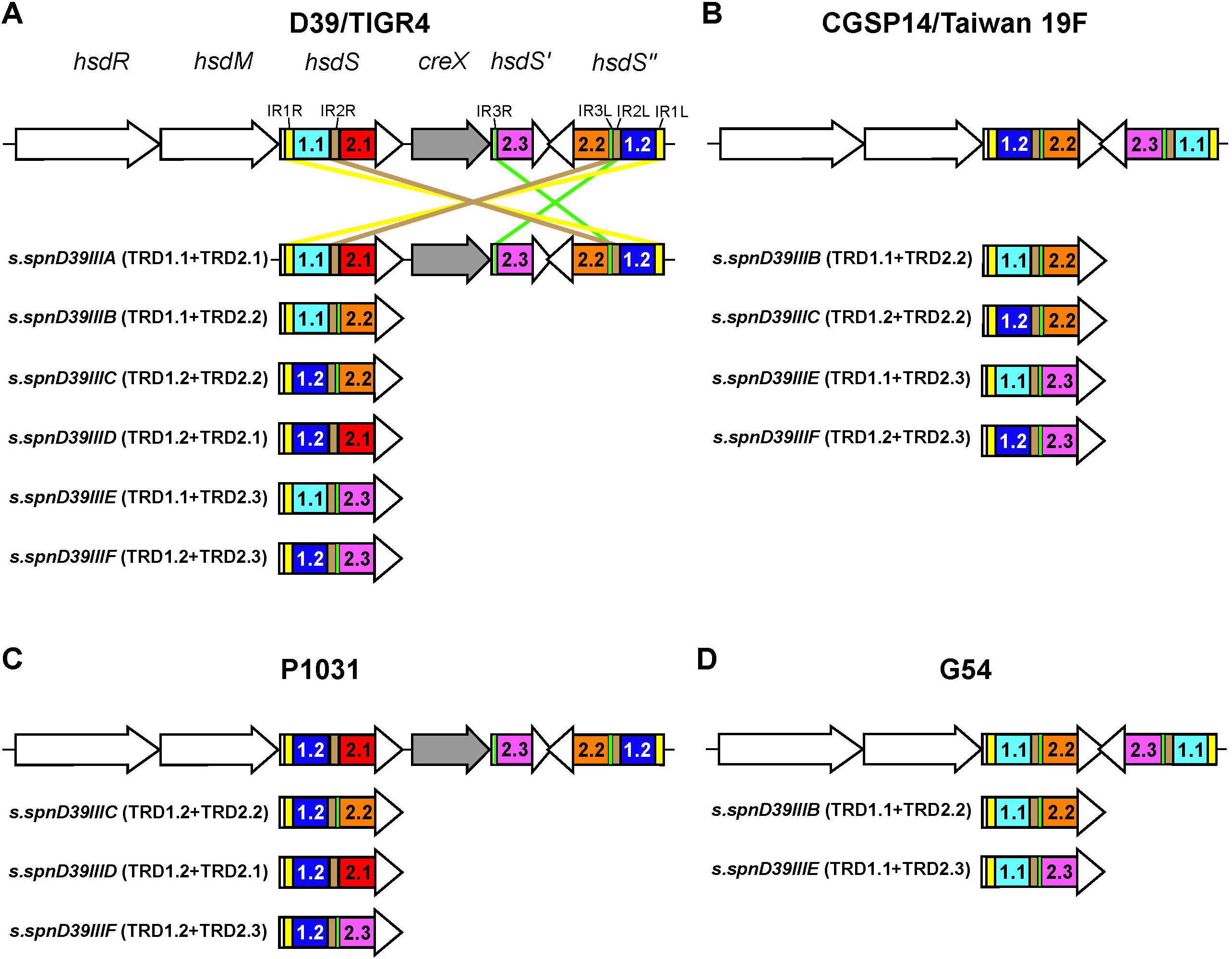
Different major forms of the SpnIII system in surveyed genomes. The SpnIII system was examined in NCBI fully annotated *S. pneumoniae* genomes. **A)** The ‘full’ six-way switch as first described in strains D39 and TIGR4. **B)** A simpler four-way switch as seen in strain CGSP14 and Taiwan 19F. **C)** A three-way switch seen in strain P1031. **D)** A two-way switch seen in strain G54. Further details can be found in **Supplementary Table 1**.

## Results

### Examination of fully annotated pneumococcal genomes reveals not all strains encode a six-way switch

The SpnIII system has been described as a six-way switch containing five characterised TRDs and a CreX recombinase [18]. We surveyed fully annotated *S. pneumoniae* genomes (82 strains total) in NCBI Genbank for the presence and conservation of the ‘full’ SpnIII system described previously [18]. **Figure 1** illustrates examples of the diversity of the SpnIII system. **Figure 1A** is the full six-way switch as seen in strains D39 and TIGR4 [18, 19], and is found in 68.3% of strains examined (56 strains). 8.5% of strains (7 strains), such as Taiwan 19F, have lost a TRD, and consequently this system only results in four unique HsdS proteins; a four-way switch producing alleles B, C, E and F (**Figure 1B**). 18.3% of strains (15 strains) encode the same TRD more than once, such as strain P1031, that encodes TRD 1.2 in both the *hsdS* and *hsdS*” loci (**Figure 1C**), producing a three-way switch, in this case, between alleles C, D and F. There are then combinations of these two factors – loss of TRDs and a duplication of one TRD, resulting in the two-way switch as seen in strain G54, which can only produce alleles B and E (**Figure 1D**). This arrangement was observed in 2.4% of surveyed genomes (2 strains). Another aspect we observed was the loss of the CreX recombinase in multiple strains (**Figure 1B, D**). The loss of this recombinase has been observed to decrease the rate of switching, but not completely prevent it [20]. Interestingly, 6.1% of strains had a SpnIII system which could not be placed into the above-described groups due to the loss or gain of identifiable features (5 strains). For example, strain 4041STDY6836167 (a serotype 9V, sequence type [ST] 156 isolate) contained a SpnIII system with seven *hsdS* regions, and strain HU-OH contained a SpnIII system with only two *hsdS* regions, one of which had <50% identity to known *hsdS* sequences (**Supplementary Table 1A**). 6.1% of genomes were also found to have two SpnIII systems. The effects of these uncharacterised TRDs, and significance of not encoding a functional SpnIII phasevarion are outside the scope of this work. All data are presented in **Supplementary Tables 1A and 1B**.

### Expression of different HsdS alleles of the full SpnIII phasevarion result in diverse, clinically relevant phenotypes

To compare clinically relevant phenotypes of the SpnIII phasevarion in strain D39, we used our six ‘locked’ D39 mutants expressing only one of the SpnIII alleles [18]. Adherence to (**Figure 2A**) and invasion of (**Figure 2B**) human lung A549 cells was investigated. Adherence was also examined using differentiated human nasal airway epithelial (HNAE) cells (**Figure 2C**). These cells, are polarised, and produce mucous. Expression of SpnIII allele B resulted in significantly more adherence to and invasion of human airway cells compared to both WT and the five other locked strains. Similar results were found using the polarised airway cell model – expression of SpnIII allele B resulted in significantly more adherence than SpnIII allele A and SpnIII allele C. When studying biofilm formation, a trait particularly important in middle ear infections, strain D39 expressing either SpnIII allele B or C showed significantly greater biofilm mass than either WT or the four other locked strains (A, D, E, F) (**Figure 2D**). Strains expressing alleles A and E showed the lowest level of biofilm formation under the conditions tested.

**Figure 2.**
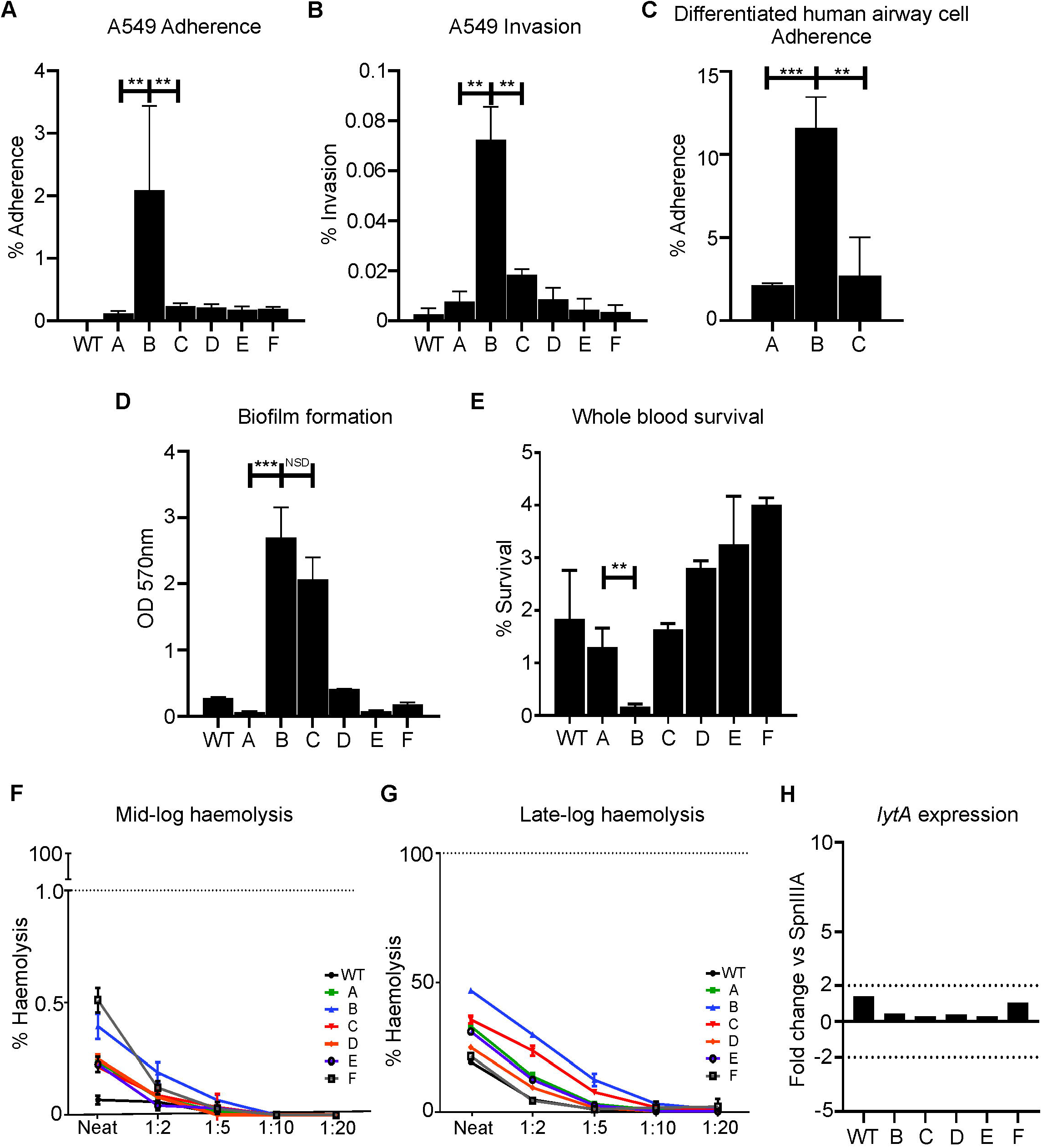
Clinically relevant traits of the SpnIII system using strain D39. **A)** Adherence of strains expressing SpnIII alleles to A549 cells. Values represent percent (%) of inoculum that was adherent. **B)** Invasion assay measuring invasive ability of strains expressing SpnIII alleles. Values represent percent of inoculum that invaded A549 cells. SpnIIIB had significantly better invasive ability compared to all other alleles. **C)** Adherence to differentiated human airway epithelial cells. Values represent percent of inoculum that adhered to cells. SpnIIIB is significantly more adherent vs A and C. **D)** Static biofilm formation was assessed via crystal violet absorbance assay. SpnIIIB and C form a significantly denser static biofilm compared to strains expressing other alleles. **E)** Ability of SpnIII alleles A-F assessed for their survival in whole human blood (group O+). SpnIIIB showed attenuated (~0%) survival, whereas SpnIIIE and F showed greatest mean survival (~3-4%). Ability of SpnIIIA-F alleles (strain D39) to lyse Human erythrocytes (group O+) measured at **F)** mid log (OD O.5) and **G)** late log (OD 1.0) growth phases. Cells expressing SpnIII allele B had the highest consistent haemolytic ability compared to other locked alleles and WT *S. pneumoniae* D39. Evaluating *lytA* expression in late log samples (**H**) showed that no significant difference between locked SpnIII alleles A-F or WT D39. This indicates differences in haemolytic ability seen in **F)** and **G)** are independent of *lytA* expression. MOI 100:1. See **Supplementary Figure 1** for challenge ratios. \**P*<0.05, \*\**P*<0.01, \*\*\**P*<0.001.

### SpnIII phase variation affects survival in blood, and haemolytic activity

All locked strains and WT exhibited 1-5% survival in human blood except the strain expressing SpnIII allele B, which was unable to survive at all in whole human blood (~0 CFU at 2h; **Figure 2E**). The haemolytic activity of each locked SpnIII strain and the WT strain was also assessed, using purified human erythrocytes (O+) as the model. The strain expressing SpnIII allele B resulted in the highest rate of haemolysis (**Figure 2F, 2G**). As haemolysis has been associated with both auto- and passive-lysis of pneumococci, we examined expression of the major autolysis associated gene – *lytA. lytA* expression was compared in each of the WT and locked strains during log-phase growth. A lower than 2-fold difference of *lytA* expression was found between all locked strains (**Figure 2H**), suggesting the differences in haemolysis are due to passive lysis, not *lytA*-mediated autolysis. All strains also expressed pneumolysin, the major toxin produced by pneumococci at similar levels, indicating that rates of haemolysis were affected by unknown factors regulated by the SpnIII phasevarion.

### The SpnIII phasevarion impacts expression of conserved protein antigens

To study the extent of SpnIII-mediated gene expression changes on conserved protein antigens, we studied expression of ten conserved pneumococcal surface proteins in prototype pneumococcal strains D39 (serotype 2) and TIGR4 (serotype 4). We used both WT and the six locked SpnIII variants in both strains D39 [16] and TIGR4 [17]. We chose ten conserved proteins that have been investigated as vaccine candidates: CpbA, GlpO, NanB, NanA, PhtD, PiuA, Ply, PsaA and PspA. Standardised lysate loads were assessed by Coomassie staining (**Supplementary Figure 2B**). Differences in protein level were observed for multiple surface proteins in both strains, (**Figure 3A, B),** with little correlation between specific SpnIII alleles and the strain being examined, indicating that there are both SpnIII phasevarion and strain-specific influences for expression of many of these conserved proteins. To investigate further, we carried out RT-pPCR using mRNA prepared from the same cultures as those used to prepare cell lysates for Western blots (**Figure 3C, 3D; full data in Supplementary Table 2A, 2B**), Significant (>2) fold differences were found in expression of vaccine candidates between the locked and WT strains, in both D39 and TIGR4. Gene expression of all vaccine candidates varied between locked strains. There was little correlation between Western blot differences and RT-qPCR expression data. To probe these differences further, whole-cell ELISAs were performed to examine the expression of targets that showed differential expression between Western blot and RT-qPCR (PiuA and PsaA) (**Supplementary Figure 2C**). We further quantified Western blot results (**Supplementary Figure 2C**) to evaluate fold difference of banding intensity, and for comparison to ELISAs. ELISAs showed protein expression profiles that aligned with the Western Blot, rather than the gene expression data. The inconsistency between mRNA and protein levels is not unusual, and has been seen in other studies [21]. This indicates that SpnIII-mediated gene expression changes, coupled with strain/serotype specific differences, play a complex role in the regulation of expression of these conserved protein antigens.

**Figure 3.**
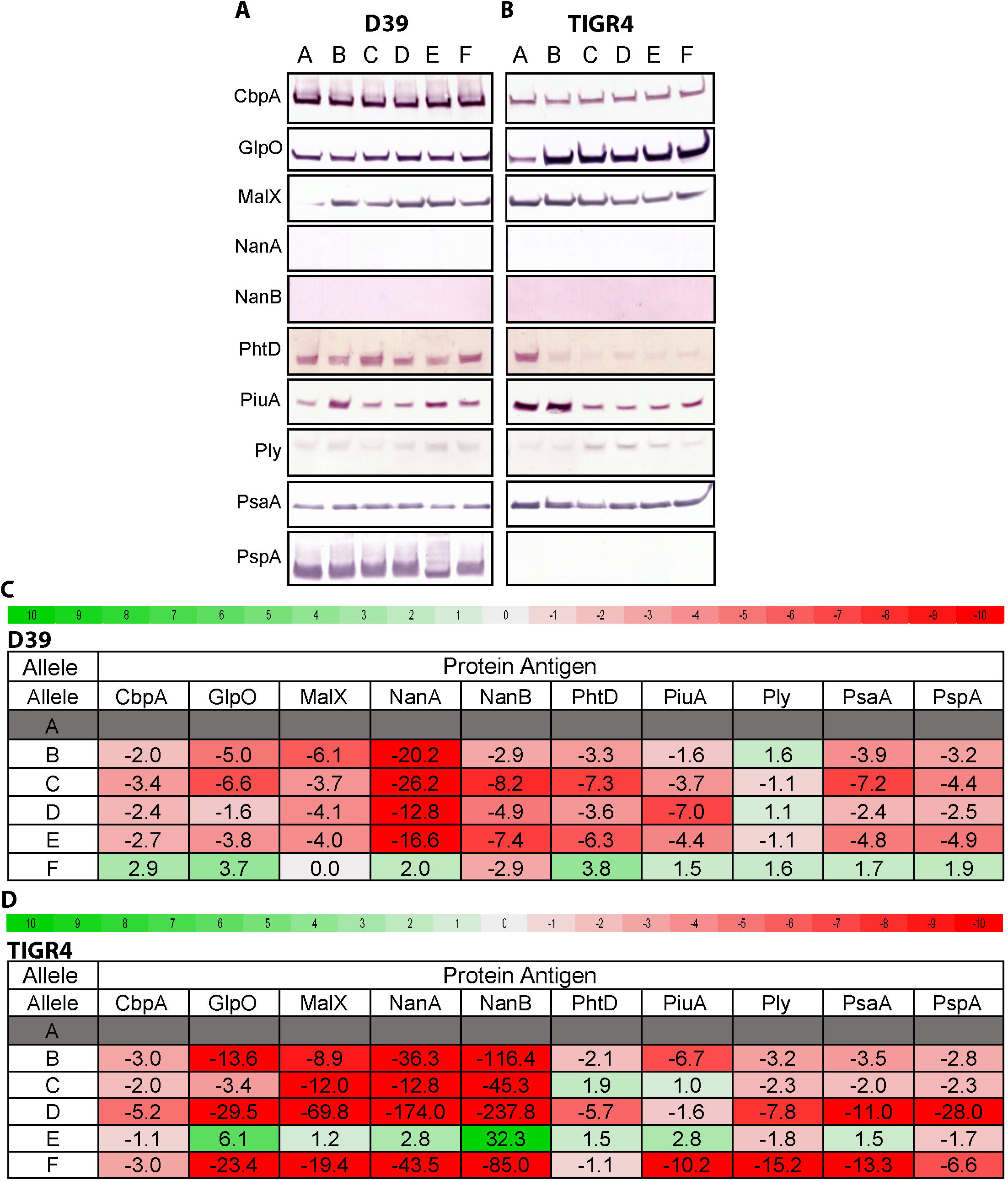
Expression differences in conserved proteinaceous vaccine candidates. Western blots examining differences in expression of protein levels between strains expressing alleles of the SpnIII system in strains **A)** D39 and **B)** TIGR4. Heatmaps of RT-qPCR expression data of concomitant expressed RNA levels of genes encoding these vaccine candidates from strains **C)** D39 and **D)** TIGR4. Red (negative fold difference) to green (positive fold difference) using SpnIII locked allele A as the baseline. Differences over >2 fold are in **bold**. All data with all locked variants as the baseline is presented in the same format in Supplementary Table 2.

### SpnIII-mediated capsule level differences are more important for pneumococcal survival than protein antigen levels

Phenotypic differences between pneumococcal strains impact complement deposition and opsonophagocytosis [22, 23]. We have shown the SpnIII variants (A-F) have different levels of protein expression of several putative protein vaccine candidates (**Figure 3**). The SpnIII system has previously been shown to influence capsule level [18]. Capsular serotype/level have also been shown to impact complement resistance and opsonophagocytosis [24]. We investigated if SpnIII phasevarion-mediated differences in both capsule quantity and protein antigen levels impacted *in vitro* opsonophagocytosis and killing by differentiated (neutrophil-like) HL-60 cells. We selected three of our D39 locked strains (locked for SpnIIIA, B and C) for their differences in capsule quantity; a D39 strain expressing SpnIII allele A has a high level of capsule relative to SpnIII allele B, with a strain expressing SpnIII allele C having an intermediate level of capsule compared to A and B [18]. SpnIII allele A (high capsule phenotype) is much more resistant to opsonophagocytic killing compared to SpnIII allele B (low capsule phenotype) and SpnIIIC (medium capsule phenotype) (**Figure 4A**). We could not achieve 50% killing of the strain locked for SpnIII allele A, even at our maximum threshold of a 400:1 MOI (neutrophil:CFU). 50% killing of SpnIII allele C was achieved with an MOI of 200:1 and SpnIII allele B with an MOI of 50:1. SpnIII allele B is over four times more susceptible to serum-independent neutrophil killing than SpnIII allele A. To investigate the effect of protein expression differences we observed in **Figure 3**, we used PiuA and CbpA anti-sera in opsonophagocytosis assays (OPKs) (**Figure 4B, 4C**). Western blots showed variable expression of PiuA commensurate with different SpnIII allele expression *(different* protein antigen level, different capsule level), whereas CbpA was equally expressed irrespective of the SpnIII allele expressed *(same* protein antigen level, different capsule level), allowing us to determine if capsule or protein antigen level was more important to survival. The large differences in antisera independent killing made comparison between locked alleles difficult. However, changes in protein antigen level impacted opsonophagocytosis less significantly than capsule level, with survival proportional to capsule amount (A>C>B).

**Figure 4.**
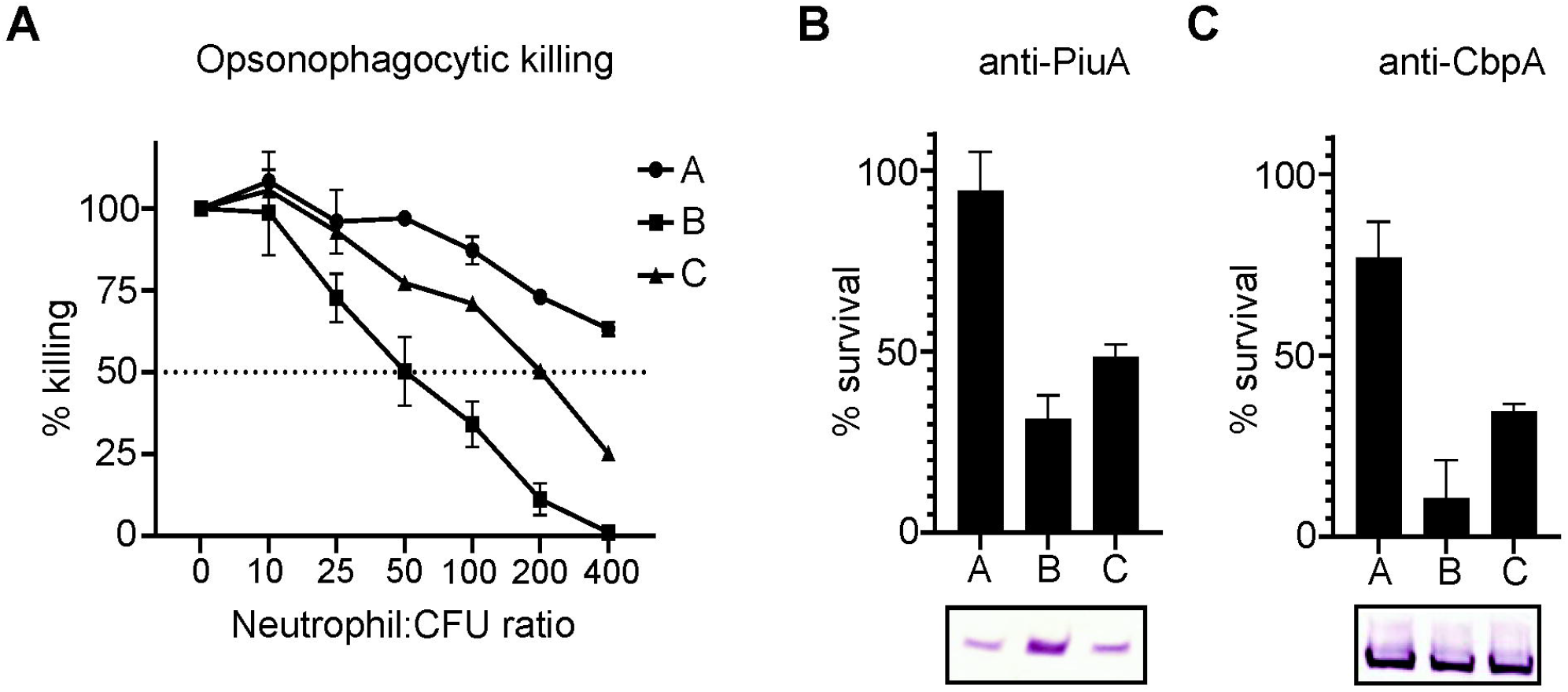
**A)** Killing by differentiated neutrophil-like HL-60 cells with 10% (v/v) complement. SpnIIIA failed to reach 50% killing at maximum neutrophil:CFU ratio (400:1) after one hour. SpnIIIB and SpnIIIC reached 50% killing at 50 and 200 neutrophil:CFU ratios, respectively. Opsonophagocytic killing assays using antisera against **B)** PiuA and **C)** CbpA at 1 in 20 dilution. Effects of antibody-mediated killing were masked by large differences in antibody-independent killing (**A**), independent of differences in anti-sera target. Western Blot showing expression of anti-sera targets in locked SpnIII alleles included for comparison. MOI for assays presented in **Supplementary Figure 1**.

## Discussion

Phasevarions offer a contingency strategy for survival in many bacteria [11, 12, 25]. In this study, we have built on our and others characterisation of the SpnIII phasevarion [18, 19]. The ‘full’ six-way SpnIII phasevarion was the most prevalent form, found in 56/82 of fully annotated *S. pneumoniae* genomes examined, indicating a key role of this system in pneumococcal pathobiology. However, we did identify variations on this system. The majority of the strains not producing a full six-way SpnIII phasevarion (24 strains) all encode systems that switch between two, three, or four SpnIII alleles (various combinations of existing alleles A-F). Five strains, such as strain 4041STDY6836167, appeared to have an SpnIII system which contained additional copies of the methyltransferase, and a total of nine *hsdS* regions (**Supplementary Table 1**), including TRDs that have not been identified before. Duplicate SpnIII systems were also observed in ~6% of genomes (5 strains). As only the six-way switch has been previously described in literature, and in only two strains (D39 and TIGR4), the impact of the truncated SpnIII phasevarions (not encoding all six HsdS alleles), the significance of duplications of the SpnIII system, and the role, if any, of newly identified TRDs, remains unclear.

We have shown that the full SpnIII phasevarion of *S. pneumoniae* produces phenotypically distinct sub-populations which have varied pathogenic phenotypes. It is therefore likely that synergy between the different phenotypes produced by the SpnIII phasevarion exists in a pneumococcal population, with each variant providing unique advantages (and disadvantages) under different conditions. These variants are likely subject to selection and counter-selection in different host niches dependent on the phenotypes they confer. There may be beneficial interactions between individual bacterial cells expressing different alleles in the pneumococcal population, with more work needed to dissect these. However, our work has begun to determine the impact of each SpnIII allele on pneumococcal phenotype on a broad population level. For example, in strain D39, SpnIII allele B produces a phenotype that is more adherent to, and invasive of, host cells, perhaps providing a mechanism for intracellular survival and persistence [26]. This property, however, comes at the expense of survival in blood, and an increased susceptibility to neutrophil-mediated killing (**Figure 4A**), with both the positive and negative phenotypes the result of much decreased capsule expression [18]. Lower capsule expression in cells expressing allele B also likely affects factors like shedding [27] and clearance [28]. We see an opposite phenotype influenced by SpnIII allele A which expresses a high amount of capsule; strains expressing this allele show a much-decreased ability to form a biofilm, and reduced adherence to host cells (**Figure 2B, 2C, 2D**), but shows high survival in blood (**Figure 2E**), and is highly resistant to neutrophil killing (**Figure 4A**). Individually, these phenotypes provide several advantages and disadvantages, but when produced in synergy they would provide a pneumococcal population with contingencies to survive in multiple host niches, and to rapidly respond to varying conditions.

We have shown that there are significant differences in haemolytic ability between the phenotypes produced by the SpnIII phasevarion during log-phase growth, independent of *lytA* expression. Haemolysis has been associated with both auto- (LytA-mediated); and passive-lysis [29–32] in *S. pneumoniae.* LytA is the major *S. pneumoniae* autolysin [29], and promotion of autolysis has been proposed to mediate release of virulence determinants such as Ply [33], and mediate *S. pneumoniae* fratricide [32]. We did not see a difference in *lytA* expression between SpnIII alleles, indicating LytA production is not the cause of haemolytic differences between strains expressing different SpnIII alleles. Therefore, the differences seen in haemolytic activity may be primarily from passive pneumococcal lysis. We also did not see a difference in pneumolysin expression (Figure 3), discounting release or level of this toxin from the differences in haemolytic ability. Manso *et al.* reported the pneumococcal capsule was significantly lower in cells expressing SpnIII alleles B and C (B<C) [18] in strain D39, and we have shown this correlates with phenotypes that can be attributed to decreased capsule. Strains expressing alleles B and C demonstrate the highest levels of haemolysis. As capsule provides structural integrity to the cell, it appears that reduced capsule, controlled by SpnIII phase-variation, produces variants that are more prone to passive lysis.

We observed stark differences in vaccine candidate expression between our ‘locked’ SpnIII strains in two diverse pneumococcal strains. Protein levels were evaluated via Western Blot (**Figure 3A, 3B, 3C**) and results were correlated with whole cell ELISA (**Supplementary Figure 2C**). There were some vaccine candidates that had relatively equal expression between alleles, such as PsaA in D39, with the largest difference of 1.5 fold between allele D and allele E (**Supplementary Figure 2C**). However, strain D39 showed different expression of MalX, PhtD and PiuA between SpnIII alleles. Quantification of the Western Blot revealed that, PiuA, for example, had a ~2.7 fold difference between D39 allele B vs allele A (**Supplementary Figure 2C**). GlpO, PhtD, Ply and PiuA protein levels showed clear expression differences in strain TIGR4. Differences seen between D39 and TIGR4 are likely due to a combination of effects, such as individual expression of different regulators/genes between the two strains, and from the proteins themselves differing between strains (e.g., D39 and TIGR4 had different families of PspA, [34, 35]). Additional factors observed to vary due to the effects of the SpnIII phasevarion, such as capsule expression [18, 19], will also affect level of cell surface proteins [28, 36]. The variation of protein antigen levels observed due to the SpnIII system may provide *S. pneumoniae* multiple contingency phenotypes, which are dynamically selected for/against depending on factors like co-colonisation and immune detection, as has been seen with phasevarions in other host-adapted pathogens [14–16, 37].

Although differences in pneumococcal genotype have been seen to impact complement deposition and opsonophagocytosis [38], capsular serotype has been observed to be the primary determinant of complement-mediated killing [24, 39]. Complicating this, the SpnIII system generates phenotypically distinct intra-strain sub-populations, which have been shown to have varied levels of pneumococcal capsule [18, 19]. We have demonstrated that even within a single strain (D39; serotype 2), the differences in capsule due to SpnIII allele switching impact neutrophil killing and likely cellular adherence and invasion (**Figure 3, 4**). Our results are in agreement with the findings of *Zangari et al.,* who recently found immunisation against conserved surface proteins induced antibody titres unable to prevent colonisation by an encapsulated strain, but were able to protect against an acapsular mutant of that same strain (that had comparable levels of antibody targets) [40]. Our findings suggest there is a particular, specialised role for a variant expressing decreased capsule level (SpnIIIB). It is possible that the protective effect seen by *Zangari et al.* when targeting conserved surface proteins in an acapsular mutant would also be effective against naturally occurring SpnIII alleles with reduced capsule (such as SpnIIIB and C in strain D39). Furthermore, targeting naturally occurring phenotypes that we have shown to be important in biofilm formation and cellular adherence and invasion may prove effective in preventing key stages of *S. pneumoniae* disease progression, such as colonisation. Several of the vaccine targets we examined (PspA, PhtD and Ply) are also known to inhibit C3 deposition [41] and interfere with complement-mediated killing. These proteins are differently expressed due to switching of the SpnIII phasevarion (**Figure 3**), demonstrating that further work is required to fully characterise the effects of the SpnIII phasevarion on killing by the host immune system.

In conclusion, we have demonstrated that clinically relevant phenotypes (**Figure 2**) conserved protein antigens (**Figure 3**), and opsonophagocytic killing (**Figure 4**) are all influenced by the SpnIII phasevarion, with many attributable to the effects of the SpnIII phasevarion on the level of capsule, correlating with previous findings [18, 19]. Significantly, we have shown that the SpnIII phasevarion has a major effect on expression of conserved protein antigens, both within individual strains, and between diverse pneumococcal strains. This is of concern to development of a universal pneumococcal vaccine based on these antigens; although these proteins are highly conserved between pneumococcal strains, their regulation appears to be profoundly complex. Understanding how expression of these antigens is regulated is critical to rational design of a universal pneumococcal vaccine, as alteration of expression of vaccine candidates could lead to a decrease in vaccine efficacy, and vaccine failure.

## Methods

### Bacterial Strains and Growth Conditions

*S. pneumoniae* strains derived from strain D39 (serotype 2) [18] or TIGR4 [19] as described previously. *S. pneumoniae* was grown in Todd Hewitt Broth supplemented with 1% Yeast (THB+Y) at 37°C. Columbia 5% Blood Agar (CBA) plates used to grow *S. pneumoniae* on solid media at 37°C 5% CO_2_ conditions.

### Adherence Assay

*S. pneumoniae* adherence and invasion was assessed against human derived A549 cells (type II pneumocytes). ~10^5^ A549 cells were seeded to each well of a flat-bottomed 24 well plate (Greiner, Germany) and allowed to settle overnight (37°C) before inoculating with *S. pneumoniae* SpnIII variants (A-F) and wild type D39 at ~10^7^ CFU per well (MOI of 100:1) in 250uL of RPMI media (Dubco) 10% foetal calf serum. MOI ratios of A549 adherence and invasion supplied in **Supplementary Figure 1A**. Plates were incubated for 45mins at 37°C 5% CO_2_ allowing adherence/invasion. For the adherence assay, wells were washed of all non-adherent *S. pneumoniae* via multiple, gentle, 1mL washes of PBS. Visual checks were performed to ensure A549s were intact, and planktonic *S. pneumoniae* removed. Wells were then detached via 250uL 0.25% Trypsin EDTA to dislodge adherent cells (5min, 37°C) before serial dilution and drop plating on CBA plates to enumerate bacterial loads. Results represent triplicate values of biological duplicates.

### Adherence to differentiated human airway epithelial cells

*S. pneumoniae* adherence was assessed using normal human nasal airway epithelial (HNAE) cells differentiated ex vivo. These primary cells were differentiated ex vivo into basal cells, ciliated cells and mucous producing cells organised in a pseudostratified epithelium that replicates the structure and nature of the human upper airway epithelium. HNAE cells were collected from healthy donors (human ethics approval GLY/01/15/HREC) and expanded using Pneumacult Ex+ (Stemcell technologies). HNAE cells were differentiated at air-liquid interface in 6.5 mm transwell with a 0.4μm polyester 156 membrane (Corning, Product No. 3470). Briefly, media was removed from HNAE cells apical side (airlift) and provided with Pneumacult ALI basal media from HNAE cells basolateral side (Stemcell technologies). HAE cells were fully differentiated and ready to use after 28 days post-airlift. Airway cells were washed twice with prewarmed phosphate buffered saline (PBS) to remove mucous (20 mins, 37°C) prior to use. The amount of total HNAE cells per transwell was enumerated, with ¼ of the total cells expected to be on the apical surface – this ¼ value was used to calculate the MOIs. Mid-log cultures of locked SpnIII alleles A, B and C from strain D39 were used to inoculate the airway cells at a surface MOI of ~5:1 in a minimal 50uL volume of media alone (Dubco). MOI ratios provided in **Supplementary Figure 1B**. CFU inputs were assessed at both initial inoculum and after one hour at 37°C to monitor changes in CFU during experimentation. Plates were incubated for 1h at 37°C with 5% (v/v) CO_2_. Wells were washed of non-adherent cells via multiple, gentle, washes with 200μL of pre-warmed PBS. Wells were then treated with 200μL of 0.25% Trypsin-EDTA to dislodge adherent bacteria (30min 37°C) before serial dilution and drop plating on Columbia-blood agar (CBA) plates to enumerate bacteria. The percentage adherence was calculated from the CFU in the inoculum. **Invasion Assay.** The Invasion Assay was identical to Adherence Assay (A549s) with the following changes: A549s were incubated with *S. pneumoniae* variants for 1h, before extracellular bacteria were killed via treatment with 50ug/mL Pen G in RPMI 10% FCS for 45min at 37°C. Wells were then treated with 250uL of 0.25% trypsin EDTA to detach A549s (5 min 37°C). Wells were then treated with 0.4% Saponin to lyse A549s (releasing invasive bacteria). Visual checks made to confirm cell lysis. Controls were in place to ensure effectiveness of Penicillin G treatment (data not shown). Surviving intracellular pneumococci were enumerated via serial dilution and drop plating on CBA plates. Results represent triplicate values of biological duplicates.

### Biofilm Formation

Log-phase *S. pneumoniae* locked alleles/wild type D39 was resuspended to OD 0.05 in THB+Y. Flat-bottomed 24 well plates (Greiner, Germany) were seeded with 1mL inoculum and grown for 24h (37°C 5% CO_2_). Wells stained and fixed with 0.5% crystal violet for 15min. Planktonic bacteria were removed via PBS washes. Remaining biomass measured by resuspending crystal violet with 95% ethanol and taking absorbance at 590nm. Results represent triplicate values of biological duplicates.

### Survival in Whole Blood

Fresh human whole blood was drawn from volunteers into heparin coated tubes by a registered nurse at Griffith University. Blood was held at 37°C, with shaking, and used within 2h of the initial bleed. Experiments were carried out in triplicate on two separate occasions. Using a 96-well flat-bottomed plate (Greiner, Germany) 200uL of whole human blood was inoculated with ~5×10^5^ CFU of strains expressing alleles A-F and wild type *S. pneumoniae* (**Supplementary Figure 1D**). Inoculum CFU were quantified by serial dilution to ensure equal loading of each well. After 90min incubation at 37°C, shaking (120rpm), with output CFU enumerated by serial dilution and drop plating. The percent survival was calculated by dividing the surviving cells in blood (output) vs inoculation dose (input). Data represents triplicate values from biological duplicates.

### Haemolysis Assay

Using 96 well round-bottomed plates (Greiner, Germany) 200uL of 10% purified human erythrocytes (Red Cross Blood Service, Australia) suspended in PBS was inoculated with 0.22u filtered minimal media of S. pneumoniae ‘locked’ strains expression SpnIII alleles A-F and D39 WT. Chemically defined media (CDM) was prepared using SILAC RPMI (no glucose, no phenol red; Life Technologies Australia) 5% Glucose and prepared as per *Minhas et al* [42]. Erythrocytes were resuspended in dilutions ranging from neat media down to a 1/100 dilution of media and PBS. Plates were incubated at 37°C, shaking at 200rpm (lid on) for 1h. Plates were removed and centrifuged at 1000rpm for 10min to pellet surviving erythrocytes. 100uL of supernatant was moved to a 96 well flat-bottomed plate (Greiner, Germany) and read at an absorbance of 540nm to detect amount of free haem and, indirectly, amount of lysis. No difference in absorbance observed between filtered minimal media and blank PBS. A PBS solution of 5% Saponin was used to produce 100% erythrocyte lysis, which was used as a standard for the amount of lysis of the *S. pneumoniae* media. Results represent triplicate values.

### Protein Extraction

To compare surface bound proteins of SpnIII variants protein was extracted from log-phase bacteria. Briefly, bacteria were grown to OD600 0.8 in TBH+Y, representing late-log phase growth. 10mL of this was pelleted and washed with THB+Y. Bacterial pellet was then resuspended in 1.5mL RIPA Buffer [43] before sonicating for 30 seconds three times, resting for 2 min between bursts. Bacteria and debris were pelleted out and supernatant (containing protein) taken. Protein lysates were stored at −20°C with 10% β-mercaptoethanol.

### Western Blotting

Protein load was standardised by visual checks on a 12% Bis-Tris Gel (**Supplementary Figure 2B**). Gels run at 150V for 45mins in Bolt MOPS Running Buffer (Invitrogen). Protein in lysates was transferred to nitrocellulose membrane at 15V for 1h. Nitrocellulose membranes were blocked with 5% (w/v) skim milk in Tris-buffered saline with 0.1% Tween-20 (TBST) by shaking overnight at 4°C. Primary mouse antibodies against *S. pneumoniae* vaccine antigens were sourced from the Paton lab (Adelaide). Primary antibodies used at 1:100-1000 in 5% skim milk TBST for 1h with shaking at room temperature. Nitrocellulose membrane was washed in TBST for 1h before adding secondary antibody (goat anti-mouse alkaline phosphatase conjugate; catalogue number A3562) (Sigma,) in 5% skim milk TBST at 1:2500 dilution. Membranes were washed for 1h in TBST, before developing for 5 – 10 mins, or overnight at room temperature with shaking. Developing solution comprised of 100mM Tris-HCL (pH 9.5), 100mM NaCl, 5mM MgCl2 with bromo-chloro-indolyl phosphate (BCIP) and nitro blue tetrazolium (NBT) used as detectors. In the case of PspA, antisera proved to be effective at detecting PspA of D39, which is classified as a family 1 PspA [44], but was infective at detecting the TIGR4 family 2 PspA [34] (evident by the lack of banding). Neither NanA nor NanB could be effectively detected in lysates after multiple attempts of varying conditions.

### *S. pneumoniae* RNA Extraction

Trizol™ protocol followed for RNA extraction of *Streptococcus spp* as per manufacturer’s instructions. Briefly, 2mL of log-phase *S. pneumoniae* at OD600 0.8 was pelleted and treated with lysozyme and mutanolysin for 30 min at 37°C before adding 1mL Trizol™ reagent and storing at −80°C. Trizol™ RNA extraction carried out as per manufacturer’s instructions (Sigma).

### Quantitative real-time PCR

Real-time PCRs were performed in triplicate using RNA isolated from ‘locked’ SpnIII alleles A-F in strain D39 and TIGR4, as described above. cDNA was synthesized using NEB Protoscript II and random hexamers (Invitrogen; 50□ng□μl^-1^) according to manufacturer instructions. Reverse transcriptase reactions lacking Protoscript II were performed as a negative control. All real-time PCR reactions were performed in a 25-μl mixture containing a 1 in 5 dilution of the cDNA preparation (5 μl), 10xSYBR Green buffer (PE Applied Biosystems) and 2 μM of each primer (See **Supplementary Table 3** for primers). Pneumococcal 16S RNA was used as a control in each quantitative PCR comparison. Amplification and detection of specific products were performed with the ABI Prism 7700 sequence-detection system (PE Applied Biosystems) with the following cycle profile: 95□°C for 1□min, followed by 40 cycles of 95□°C for 30s and 60 °C for 1□min. The data was analysed with ABI prism 7700 (version 1.7) analysis software. Relative gene expression between samples was determined using the ^ΔΔ^CT relative quantification method.

### Opsonophagocytic killing assays

HL60 cells were differentiated in DMSO (0.8%) containing M2 media (RPMI 1640, 10% FBS, 1% L-glutamine) for six days prior to use. To determine HL60 differentiation, flow cytometry analysis was carried out to determine >55% of cells expressing the maturation marker CD35 (E11, Biorad) and >12% of cells expressing the proliferation marker CD71 (DF1513, Biorad) (PMID: 32430834). Differentiated HL60s were activated in Hank’s balanced salt solution (HBSS) containing Ca^++^ and Mg^++^ prior to use. The ability of the differentiated HL60 neutrophil-like cells to kill *S. pneumoniae* D39 locked alleles A, B and C was assessed across varying concentrations of rabbit complement and neutrophils with a range of 20% complement + 4×10^5^ Neutrophils per 1×10^3^ CFU of *S. pneumoniae* down to complement and neutrophil free media. Complement + media only controls did not kill *S. pneumoniae* (data not shown). Killing was assessed after 1h at 37°C in conditions with and without mouse anti-sera. Results were expressed as percent of the surviving inoculum in the control wells for the neutrophil killing assay, and percent survival of the starting inoculum (~1000 CFU) in the opsonophagocytosis assays. Input amounts of each Locked allele can be seen in **Supplementary Figure 1C**. **Statistical Analysis.** Graphs and statistics were generated via GraphPad Prism 5.0 (GraphPad Software, La Jolla, California). Error bars represent standard deviation from mean values. Student’s *t* test was used to compare samples: p values of <0.05 (considered significant) represented by *, p value of <0.001 indicated by **, p value of <0.001 indicated by ***. Groups were considered not significantly different if p > 0.05 (no *).

## Acknowledgments

This work was supported by Australian Research Council (ARC) Discovery Project Grant DP180100976 to J.M.A., ARC Discovery Project Grant DP190102980 to C.T. and J.C.P., National Health and Medical Research Council (NHMRC) Program Grant 1071659 to J.C.P., M.P.J. and M. vI., NHMRC Investigator Grant 1174876 to J.C.P., NHMRC Principal Research Fellowship 1138466 to M.P.J, and NHMRC Ideas Grant APP1160379 to M.P. We thank Griffith University for providing ZP with a PhD scholarship. Publication costs of this work were supported by a generous donation from the Bourne Foundation, Melbourne, Australia.

## Notes

### Competing Interest Statement

The authors have declared no competing interest.

