## Supplementary Tables for "A phasevarion controls multiple virulence traits, including expression of vaccine candidates, in *Streptococcus pneumoniae*"

## A

| | SpnIII System Genetic Region | | | | | | | | | | |
|---|---|---|---|---|---|---|---|---|---|---|---|
| Strain | 1 | 2 | 3 | 4 | 5 | 6 | 7 | 8 | 9 | Methyltransferase | Type |
| 310 | 1.1> | 1.2> | PsrA> | 2.3> | <2.2 | <1.1 | | | | <M | 3 |
| 335 | 1.1> | 2.3> | <2.2 | <PsrA | <2.1 | <1.2 | | | | <M | 1 |
| 475 | 1.1> | 1.2> | <2.2 | <PsrA | <2.1 | <1.2 | | | | <M | 3 |
| 521 | 1.2> | 2.1> | PsrA> | 2.3> | <2.2 | <1.1 | | | | <M | 1 |
| 521 | 1.2> | 2.1> | PsrA> | 2.2> | <2.3 | <1.1 | | | | <M | 1 |
| 525 | 1.1> | 2.1> | PsrA> | 2.3> | <2.2 | <1.1 | | | | <M | 1 |
| 563 | 1.2> | 2.1> | PsrA> | 2.2> | <2.3 | <1.1 | | | | <M | 1 |
| 566 | 1.1> | 2.1> | PsrA> | 2.3> | ? | <1.2 | | | | <M | 1 |
| 573 | 1.2> | 2.2> | <2.3 | <PsrA | <2.1 | <1.1 | | | | <M | 1 |
| 574 | 1.2> | 2.1< | PsrA> | 2.3< | <2.2 | <1.1 | | | | <M | 1 |
| 947 | 1.1> | 2.1> | PsrA> | ? | ? | <1.1 | | | | <M | 3 |
| 4496 | 1.2> | 2.1> | PsrA> | 2.3> | <2.2 | <1.1 | | | | <M | 1 |
| 70585 | 1.1> | 2.3> | <2.2 | <1.2 | | | | | | <M | 2 |
| 11A | 1.2> | 2.1> | PsrA> | 2.3> | <2.2 | <1.1 | | | | <M | 1 |
| 180-15 | 1.2> | 2.3> | <2.2 | <PsrA | <2.1 | <1.1 | | | | <M | 1 |
| 180-2 | 1.1> | 2.2> | <2.3 | <PsrA | <2.1 | <1.2 | | | | <M | 1 |
| 19A-19339 | 1.2> | 2.3> | <2.2 | <1.2 | | | | | | <M | 4 |
| 2245STDY6178787 | 1.2> | 2.1> | PsrA> | 2.3> | 2.2> | <2.3 | <1.1 | | | <M | 5 |
| 4041STDY6583227 | 1.2> | 2.1> | PsrA> | 2.3> | <2.2 | <1.1 | | | | <M | 1 |
| 4041STDY6836166 | <2.2 | <PsrA | <2.1 | <1.2 | | | | | | <M | 5 |
| 4041STDY6836166_2 | 1.2> | 2.1> | <PsrA | 2.2> | 2.3> | <1.2 | | | | <M | 3 |
| 4041STDY6836167 | 2.2> | <2.3 | M> | 1.2> | 2.2> | <2.3 | <PsrA | <2.1 | <1.2 | <M | 5 |
| 4041STDY6836169 | 1.2> | 2.1> | PsrA> | 2.3> | <2.2 | <1.1 | | | | <M | 1 |
| 4041STDY6836169_2 | 1.1> | 2.1> | PsrA> | 2.2> | <2.3 | <1.1 | | | | <M | 3 |
| 4041STDY6836170 | 1.2> | 2.2> | <2.3 | <PsrA | <2.1 | <1.2 | | | | <M | 3 |
| 670-6B | 1.2> | 2.3> | <2.2 | <PsrA | <2.1 | <1.1 | | | | <M | 1 |
| 6A-10 | 1.2> | 2.1> | PsrA> | 2.3> | <2.2 | <1.1 | | | | <M | 1 |
| A45 | | | | | | | | | | | |
| A66 | | | | | | | | | | | |
| AP200 | 1.2> | 2.2> | <2.3 | <PsrA | <2.1 | <1.1 | | | | <M | 1 |
| ASP0581 | 1.1> | 2.3> | <2.2 | <PsrA | <2.1 | <1.2 | | | | <M | 1 |
| ATCC 49619 | 1.2> | 2.1> | PsrA> | 2.3> | <2.2 | <1.1 | | | | <M | 1 |
| ATCC 700669 | 1.1> | 2.3> | <2.2 | <PsrA | <2.1 | <1.2 | | | | <M | 1 |
| AUSMDU00010538 | 1.2> | 2.1> | PsrA> | ? | ? | <1.1 | | | | <M | 1 |
| B1900 | 1.2> | 2.2> | <2.3 | <1.1 | | | | | | <M | 2 |
| BHN97x | 1.2> | 2.1> | PsrA> | 2.3> | <2.2 | <1.2 | | | | <M | 3 |
| BVJ1JL | 1.2> | 2.2> | <2.3 | <PsrA | <2.1 | <1.2 | | | | <M | 3 |
| CGSP14 | 1.1> | 2.3> | <2.2 | <1.2 | | | | | | <M | 2 |
| D39 | 1.2> | 2.2< | <2.3 | <PsrA | <2.1 | <1.1 | | | | <M | 1 |
| EF3030 | ? | 2.1> | PsrA> | ? | ? | <1.1 | | | | <M | 1 |
| FDAARGOS-1508 | 1.2> | 2.1> | PsrA> | 2.2> | <2.3 | <1.1 | | | | <M | 1 |
| G54 | 1.1> | 2.2> | <2.3 | <1.1 | | | | | | <M | 4 |
| gamPNI0373 | 1.2> | 2.1> | PsrA> | 2.3> | <2.2 | <1.2 | | | | <M | 3 |
| GPSC10 | 1.2> | 2.1> | ? | ? | | | | | | <M | 2 |
| GPSC10_2 | 1.2> | 2.2> | <2.3 | <PsrA | <2.1 | <1.1 | | | | <M | 1 |
| GPSC18 | 2.2> | <2.3 | ? | <PsrA | <2.1 | <1.1 | | | | <M | 1 |
| HKU1-14 | 1.2> | 2.2> | <2.3 | <PsrA | <2.1 | <1.2 | | | | <M | 3 |
| Hungary19A-6 | 1.2> | 2.1> | PsrA> | 2.3> | <2.2 | <1.1 | | | | <M | 1 |
| HU-OH | ? | <1.1 | | | | | | | | <M | 5 |
| INV104 | 1.2> | 2.3> | <2.2 | <1.1 | | | | | | <M | 2 |
| INV200 | ? | 2.1> | PsrA> | 2.3> | <2.2 | <1.1 | | | | <M | 1 |
| JJA | 1.2> | 2.3> | <2.2 | <PsrA | <2.1 | <1.1 | | | | <M | 1 |
| KK0981 | 1.2> | 2.1> | PsrA> | 2.3> | <1.2 | <1.1 | | | | <M | 3 |
| M16808 | 1.1> | 2.1> | PsrA> | 2.2> | <2.3 | <1.2 | | | | <M | 1 |
| M23734 | 1.2> | 2.1> | PsrA> | 2.3> | <2.2 | <1.2 | | | | <M | 3 |
| M26365 | 1.2> | 2.2> | <2.3 | <PsrA | <2.1 | <1.1 | | | | <M | 1 |
| M26368 | 1.1> | 2.1> | PsrA> | 2.3> | <2.2 | <1.2 | | | | <M | 1 |
| MDRSPN001 | 1.1> | 2.2> | <2.3 | <PsrA | <2.1 | <1.2 | | | | <M | 1 |
| NCTC11902 | 1.2> | 2.1> | PsrA> | 2.3> | <2.2 | <1.1 | | | | <M | 1 |
| NCTC12977 | 1.1> | 2.1> | PsrA> | 2.2> | <2.3 | <1.1 | | | | <M | 1 |
| NCTC12977_2 | 1.1> | 2.1> | PsrA> | 2.2> | <2.3 | <1.2 | | | | <M | 1 |
| NCTC13276 | 1.2> | 2.2> | <2.3 | <PsrA | <2.1 | <1.1 | | | | <M | 1 |
| NCTC7465 | 1.2> | >2.3 | <2.2 | <PsrA | <2.1 | <1.1 | | | | <M | 1 |
| NCTC7465_2 | 1.2> | 2.1> | <PsrA | 2.3> | <2.2 | <1.1 | | | | <M | 1 |
| NCTC7466 | 1.2> | 2.2> | <2.3 | <PsrA | <2.1 | <1.1 | | | | <M | 1 |
| NT-110-58 | 1.1> | 2.1> | PsrA> | 2.2> | <2.3 | <1.2 | | | | <M | 1 |
| NU83127 | 1.1> | 2.1> | <PsrA | 2.2> | <2.3 | <1.1 | | | | <M | 3 |
| OXC141 | 1.2> | 2.3> | <2.2 | <PsrA | <2.1 | <1.1 | | | | <M | 1 |
| P1031 | 1.2> | 2.1> | PsrA> | 2.3> | <2.2 | <1.2 | | | | <M | 3 |
| PZ900701590 | 1.1> | 2.1> | PsrA> | 2.3> | <2.2 | <1.1 | | | | <M | 3 |
| R36A | 1.1> | 2.2> | <2.3 | <PsrA | <2.1 | <1.2 | | | | <M | 1 |
| R6 | 1.2> | 2.2> | <2.3 | <PsrA | <2.1 | <1.1 | | | | <M | 1 |
| R6CIB17 | 1.2> | 2.2> | <2.3 | <PsrA | <2.1 | <1.1 | | | | <M | 1 |
| Rx1 | 1.2> | 2.1> | PsrA> | 2.3> | <2.2 | <1.1 | | | | <M | 1 |
| SCAID PHRX1-2021 | 1.2> | 2.1> | PsrA> | 2.3> | <2.2 | <1.1 | | | | <M | 1 |
| SNP034156 | 1.2> | 2.1> | PsrA> | 2.2> | <2.3 | <1.1 | | | | <M | 1 |
| SP49 | 1.2> | 2.1> | PsrA> | 2.2> | <2.3 | <1.1 | | | | <M | 1 |
| SP61 | 1.2> | 2.1> | PsrA> | 2.2> | <2.3 | <1.1 | | | | <M | 1 |
| SP64 | 1.2> | 2.1> | PsrA> | 2.2> | <2.3 | <1.1 | | | | <M | 1 |
| SPN XDR SMC1710-32 | <PsrA | <2.1 | <1.1 | | | | | | | <M | 5 |
| SPN032672 | 1.2> | 2.3> | <2.2 | <1.1 | | | | | | <M | 2 |
| ST556 | 1.2> | 2.1> | PsrA> | 2.2> | <2.3 | <1.1 | | | | <M | 1 |
| SWU02 | 1.1> | 2.1> | PsrA> | 2.2> | <2.3 | <1.2 | | | | <M | 1 |
| Taiwan19F-14 | 1.2> | 2.2> | <2.3 | <1.1 | | | | | | <M | 2 |
| TCH8431/19A | 1.2> | 2.1> | PsrA> | 2.3> | <2.2 | <1.1 | | | | <M | 1 |
| TIGR4; ATCC BAA-334 | 1.2> | 2.1> | PsrA> | 2.3> | <2.2 | <1.1 | | | | <M | 1 |
| Xen35 | 1.2> | 2.1> | PsrA> | 2.3> | <2.2 | <1.1 | | | | <M | 1 |

**B**

| SpnIII Type | Prevalence in Genomes |
|---|---|
| 1 | 68.3% |
| 2 | 8.5% |
| 3 | 18.3% |
| 4 | 2.4% |
| 5 | 6.1% |
| Two SpnIII Systems | 6.1% |
| Absent | 2.4% |

**Supplementary Table 1.** SpnIII systems in fully annotated NCBI *S. pneumoniae* genomes. **A)** Orientation and genetic features of SpnIII systems. Direction of each feature indicated with either > or <. *hsdS* sequences 1.1, 1.2, 2.1, 2.2 and 2.3 included. Red highlighted sections have low (<80%) sequence identity to known *hsdS* sequences. "?" indicates a *hsdS* region with an identity <50% to the known *hsdS* sequences. Duplications of features are indicated in Green. The CreX recombinase – Pneumococcal site-specific recombinase A (PsrA) included in **Bold**. The sequence for the associated Methyltransferase (M) was also included with orientation. The prevalence of each major type of SpnIII system found in genomes can be seen in **B)**. Type 1 is the 'traditional' six-way switch as seen in **Figure 1A**. Type 2, seen in **Figure 1B**, is a four-way switch. Type 3 is a three-way switch as per **Figure 1C**. Type 4 is a two-way switch, as per **Figure 1D**. Type 5 were instances that did not fall into the previous four categories.

| A | Protein Antigen Expression in D39 | | | | | | | | | |
|---|---|---|---|---|---|---|---|---|---|---|
| **Allele** | **CbpA** | **GlpO** | **MalX** | **NanA** | **NanB** | **PhtD** | **PiuA** | **Ply** | **PsaA** | **PspA** |
| A | | | | | | | | | | |
| B | -2.0 | -5.0 | -6.1 | -20.2 | -2.9 | -3.3 | -1.6 | 1.6 | -3.9 | -3.2 |
| C | -3.4 | -6.6 | -3.7 | -26.2 | -8.2 | -7.3 | -3.7 | -1.1 | -7.2 | -4.4 |
| D | -2.4 | -1.6 | -4.1 | -12.8 | -4.9 | -3.6 | -7.0 | 1.1 | -2.4 | -2.5 |
| E | -2.7 | -3.8 | -4.0 | -16.6 | -7.4 | -6.3 | -4.4 | -1.0 | -4.7 | -4.9 |
| F | 2.0 | 2.4 | -1.1 | 1.4 | -3.5 | 1.2 | 1.1 | 1.5 | 1.3 | 1.3 |

| Allele | CbpA | GlpO | MalX | NanA | NanB | PhtD | PiuA | Ply | PsaA | PspA |
|---|---|---|---|---|---|---|---|---|---|---|
| A | 2.0 | 5.0 | 6.1 | 20.2 | 2.9 | 3.3 | 1.6 | -1.6 | 3.9 | 3.2 |
| B | | | | | | | | | | |
| C | -1.7 | -1.3 | 1.6 | -1.3 | -2.8 | -2.2 | -2.4 | -1.8 | -1.8 | -1.4 |
| D | -1.2 | 3.0 | 1.5 | 1.6 | -1.7 | -1.1 | -4.5 | -1.5 | 1.7 | 1.3 |
| E | -1.4 | 1.3 | 1.5 | 1.2 | -2.5 | -1.9 | -2.8 | -1.7 | -1.2 | -1.5 |
| F | 4.0 | 11.8 | 5.6 | 27.8 | -1.2 | 4.0 | 1.8 | -1.1 | 5.0 | 4.3 |

| Allele | CbpA | GlpO | MalX | NanA | NanB | PhtD | PiuA | Ply | PsaA | PspA |
|---|---|---|---|---|---|---|---|---|---|---|
| A | 3.4 | 6.6 | 3.7 | 26.2 | 8.2 | 7.3 | 3.7 | 1.1 | 7.2 | 4.4 |
| B | 1.7 | 1.3 | -1.6 | 1.3 | 2.8 | 2.2 | 2.4 | 1.8 | 1.8 | 1.4 |
| C | | | | | | | | | | |
| D | 1.4 | 4.0 | -1.1 | 2.1 | 1.7 | 2.0 | -1.9 | 1.2 | 3.0 | 1.8 |
| E | 1.3 | 1.8 | -1.1 | 1.6 | 1.1 | 1.2 | -1.2 | 1.1 | 1.5 | -1.1 |
| F | 6.8 | 15.8 | 3.4 | 36.0 | 2.4 | 8.7 | 4.2 | 1.7 | 9.1 | 5.9 |

| Allele | CbpA | GlpO | MalX | NanA | NanB | PhtD | PiuA | Ply | PsaA | PspA |
|---|---|---|---|---|---|---|---|---|---|---|
| A | 2.4 | 1.6 | 4.1 | 12.8 | 4.9 | 3.6 | 7.0 | -1.1 | 2.4 | 2.5 |
| B | 1.2 | -3.0 | -1.5 | -1.6 | 1.7 | 1.1 | 4.5 | 1.5 | -1.7 | -1.3 |
| C | -1.4 | -4.0 | 1.1 | -2.1 | -1.7 | -2.0 | 1.9 | -1.2 | -3.0 | -1.8 |
| D | | | | | | | | | | |
| E | -1.1 | -2.3 | 1.0 | -1.3 | -1.5 | -1.7 | 1.6 | -1.1 | -2.0 | -2.0 |
| F | 4.8 | 3.9 | 3.8 | 17.6 | 1.4 | 4.3 | 8.1 | 1.4 | 3.0 | 3.3 |

| Allele | CbpA | GlpO | MalX | NanA | NanB | PhtD | PiuA | Ply | PsaA | PspA |
|---|---|---|---|---|---|---|---|---|---|---|
| A | 2.7 | 3.8 | 4.0 | 16.6 | 7.4 | 6.3 | 4.4 | 1.0 | 4.7 | 4.9 |
| B | 1.4 | -1.3 | -1.5 | -1.2 | 2.5 | 1.9 | 2.8 | 1.7 | 1.2 | 1.5 |
| C | -1.3 | -1.8 | 1.1 | -1.6 | -1.1 | -1.2 | 1.2 | -1.1 | -1.5 | 1.1 |
| D | 1.1 | 2.3 | 1.0 | 1.3 | 1.5 | 1.7 | -1.6 | 1.1 | 2.0 | 2.0 |
| E | | | | | | | | | | |
| F | 5.3 | 9.0 | 3.7 | 22.8 | 2.1 | 7.5 | 5.0 | 1.6 | 6.0 | 6.6 |

| Allele | CbpA | GlpO | MalX | NanA | NanB | PhtD | PiuA | Ply | PsaA | PspA |
|---|---|---|---|---|---|---|---|---|---|---|
| A | -2.0 | -2.4 | 1.1 | -1.4 | 3.5 | -1.2 | -1.1 | -1.5 | -1.3 | -1.3 |
| B | -4.0 | -11.8 | -5.6 | -27.8 | 1.2 | -4.0 | -1.8 | 1.1 | -5.0 | -4.3 |
| C | -6.8 | -15.8 | -3.4 | -36.0 | -2.4 | -8.7 | -4.2 | -1.7 | -9.1 | -5.9 |
| D | -4.8 | -3.9 | -3.8 | -17.6 | -1.4 | -4.3 | -8.1 | -1.4 | -3.0 | -3.3 |
| E | -5.3 | -9.0 | -3.7 | -22.8 | -2.1 | -7.5 | -5.0 | -1.6 | -6.0 | -6.6 |
| F | | | | | | | | | | |

| B | Protein Antigen Expression in TIGR4 | | | | | | | | | |
|---|---|---|---|---|---|---|---|---|---|---|
| **Allele** | **CbpA** | **GlpO** | **MalX** | **NanA** | **NanB** | **PhtD** | **PiuA** | **Ply** | **PsaA** | **PspA** |
| A | | | | | | | | | | |
| B | 3.0 | 13.6 | 8.9 | 36.3 | 116.4 | 2.0 | 6.7 | 3.2 | 3.5 | 2.7 |
| C | 2.0 | 3.4 | 12.0 | 12.7 | 45.3 | -1.9 | -1.0 | 2.3 | 2.0 | 2.3 |
| D | 5.2 | 29.5 | 69.8 | 174.0 | 246.8 | 5.7 | 1.6 | 7.8 | 11.0 | 27.9 |
| E | 1.1 | -6.1 | -1.2 | -2.8 | -32.3 | -1.5 | -2.8 | 1.8 | -1.5 | 1.7 |
| F | 3.0 | 23.4 | 19.4 | 43.5 | 84.9 | 1.1 | 10.2 | 15.2 | 13.3 | 6.6 |
| **Allele** | **CbpA** | **GlpO** | **MalX** | **NanA** | **NanB** | **PhtD** | **PiuA** | **Ply** | **PsaA** | **PspA** |
| A | -3.0 | -13.6 | -8.9 | -36.3 | -116.4 | -1.7 | -6.7 | -3.2 | -3.5 | -2.7 |
| B | | | | | | | | | | |
| C | -1.5 | -4.0 | 1.3 | -2.8 | -2.6 | -3.2 | -6.7 | -1.4 | -1.7 | -1.2 |
| D | 1.7 | 2.2 | 7.8 | 4.8 | 2.1 | 3.3 | -4.3 | 2.4 | 3.1 | 10.2 |
| E | -2.7 | -82.8 | -10.3 | -102.5 | -3762.8 | -2.6 | -18.9 | -1.8 | -5.1 | -1.6 |
| F | 1.0 | 1.7 | 2.2 | 1.2 | -1.4 | -1.5 | 1.5 | 4.7 | 3.8 | 2.4 |
| **Allele** | **CbpA** | **GlpO** | **MalX** | **NanA** | **NanB** | **PhtD** | **PiuA** | **Ply** | **PsaA** | **PspA** |
| A | -2.0 | -3.4 | -12.0 | -12.7 | -45.3 | 3.5 | 1.0 | -2.3 | -2.0 | -2.3 |
| B | 1.5 | 4.0 | -1.3 | 2.8 | 2.6 | 7.1 | 6.7 | 1.4 | 1.7 | 1.2 |
| C | | | | | | | | | | |
| D | 2.6 | 8.7 | 5.8 | 13.6 | 5.5 | 19.7 | 1.6 | 3.4 | 5.4 | 12.3 |
| E | -1.8 | -20.8 | -13.8 | -36.0 | -1463.5 | 2.3 | -2.8 | -1.3 | -2.9 | -1.3 |
| F | 1.5 | 6.9 | 1.6 | 3.4 | 1.9 | 3.9 | 10.2 | 6.7 | 6.5 | 2.9 |
| **Allele** | **CbpA** | **GlpO** | **MalX** | **NanA** | **NanB** | **PhtD** | **PiuA** | **Ply** | **PsaA** | **PspA** |
| A | -5.2 | -29.5 | -69.8 | -174.0 | -246.8 | -5.7 | -1.6 | -7.8 | -11.0 | -27.9 |
| B | -1.7 | -2.2 | -7.8 | -4.8 | -2.1 | -2.8 | 4.3 | -2.4 | -3.1 | -10.2 |
| C | -2.6 | -8.7 | -5.8 | -13.6 | -5.5 | -10.6 | -1.6 | -3.4 | -5.4 | -12.3 |
| D | | | | | | | | | | |
| E | -4.7 | -179.7 | -80.6 | -491.0 | -7981.1 | -8.5 | -4.4 | -4.4 | -16.0 | -16.0 |
| F | -1.7 | -1.3 | -3.6 | -4.0 | -2.9 | -5.0 | 6.5 | 2.0 | 1.2 | -4.2 |
| **Allele** | **CbpA** | **GlpO** | **MalX** | **NanA** | **NanB** | **PhtD** | **PiuA** | **Ply** | **PsaA** | **PspA** |
| A | -1.1 | 6.1 | 1.2 | 2.8 | 32.3 | 1.5 | 2.8 | -1.8 | 1.5 | -1.7 |
| B | 2.7 | 82.8 | 10.3 | 102.5 | 3762.8 | 3.1 | 18.9 | 1.8 | 5.1 | 1.6 |
| C | 1.8 | 20.8 | 13.8 | 36.0 | 1463.5 | -1.3 | 2.8 | 1.3 | 2.9 | 1.3 |
| D | 4.7 | 179.7 | 80.6 | 491.0 | 7981.1 | 8.5 | 4.4 | 4.4 | 16.0 | 16.0 |
| E | | | | | | | | | | |
| F | 2.7 | 143.0 | 22.4 | 122.9 | 2747.0 | 1.7 | 28.7 | 8.6 | 19.3 | 3.8 |
| **Allele** | **CbpA** | **GlpO** | **MalX** | **NanA** | **NanB** | **PhtD** | **PiuA** | **Ply** | **PsaA** | **PspA** |
| A | -3.0 | -23.4 | -19.4 | -43.5 | -84.9 | -1.1 | -10.2 | -15.2 | -13.3 | -6.6 |
| B | -1.0 | -1.7 | -2.2 | -1.2 | 1.4 | 1.8 | -1.5 | -4.7 | -3.8 | -2.4 |
| C | -1.5 | -6.9 | -1.6 | -3.4 | -1.9 | -2.1 | -10.2 | -6.7 | -6.5 | -2.9 |
| D | 1.7 | 1.3 | 3.6 | 4.0 | 2.9 | 5.0 | -6.5 | -2.0 | -1.2 | 4.2 |
| E | -2.7 | -143.0 | -22.4 | -122.9 | -2747.0 | -1.7 | -28.7 | -8.6 | -19.3 | -3.8 |
| F | | | | | | | | | | |

**Supplementary Table 2.** Putative vaccine target RTqPCR. Heatmap of putative vaccine target gene expression across strains **A)** D39 and **B)** TIGR4 expressing individual SpnDIII alleles ranging from red

(- fold difference) to green (+ fold difference). Gene expression values from locked alleles in grey have been used as a baseline against other alleles.

| Primer Name | Forward/Reverse | Sequence (5' - 3') |
|---|---|---|
| CbpA_RT_F | F | CAAGGTAAACCAAAGGGGCG |
| CbpA_RT_R | R | TCAGGGATGAGCTTGGAAGAG |
| GlpO_RT_F | F | CATTGCCAACCACGTGAAGG |
| GlpO_RT_R | R | AACCAGACGGGCCTTGATTT |
| MalX_RT_F | F | TACGCATTCGCTGGTGAAGA |
| MalX_RT_R | R | AGCGTCTTTACCGTTTTGGC |
| NanA_RT_F | F | ATGACGACGGGAAGACATGG |
| NanA_RT_R | R | TTGTGAGGCCCATTCCGAAG |
| NanB_RT_F | F | TCTAGAAGTGGCCGTAAGGGA |
| NanB_RT_R | R | AGGCATAACCATACGAAGGCAA |
| PhtD_RT_F | F | AGCTGTTCGAAAAGTAGGCGA |
| PhtD_RT_R | R | ACTTTCCTGCTTGGCCAGTT |
| Ply_RT_F | F | CCCACTCTTCTTGCGGTTGA |
| Ply_RT_R | R | TCCGCGAACACTTGAATTGC |
| PspA_RT_R | F | CGCTCCTCAAGCTAAAATCGC |
| PspA_RT_R | R | GAAGAGGAGCACGGAAACCT |
| PiuA_RT_F | F | CCGAAGGCACTTGCTAAGGA |
| PiuA_RT_R | R | TCGCTTCACCACGTACACAA |
| PsaA_RT_F | F | CAGCGACGGCGTTGATGTTA |
| PsaA_RT_R | R | TTGGCGCTCAATTGTTTGGC |
| lytA_RT_F | F | CGGTTGGAATGCTGAGACCT |
| lytA_RT_R | R | GGCAAACCTGCTTCATCTGC |

**Supplementary Table 3.** RTqPCR Primers.
