## Supplementary Figure 1 for "A phasevarion controls multiple virulence traits, including expression of vaccine candidates, in *Streptococcus pneumoniae*"

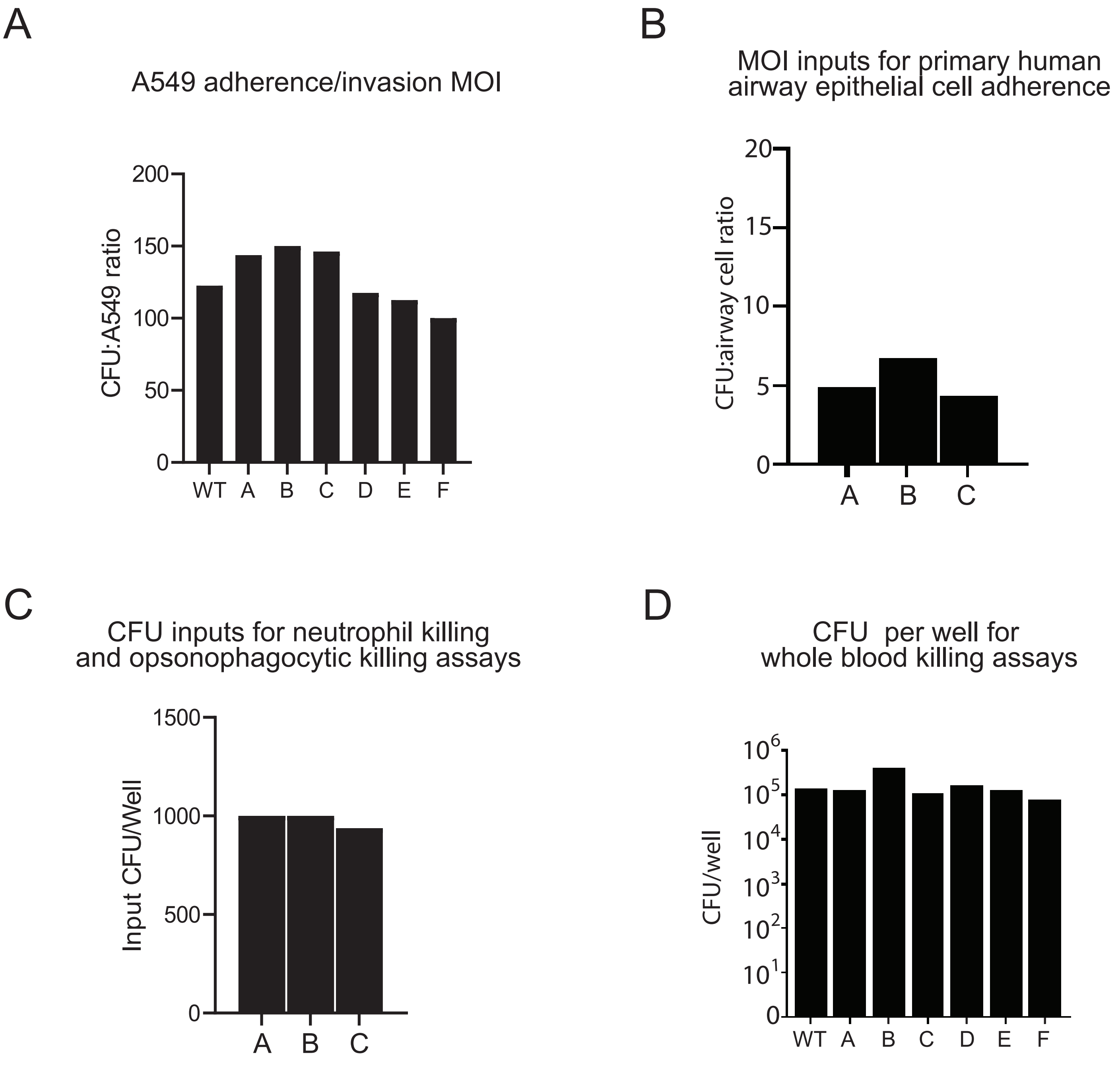


**Supplementary Figure 1**. Balanced *S. pneumoniae* D39 SpnIII locked allele inputs. Input colony forming unit (CFU) loads were standardised for experiments. A multiplicity of infection (MOI) of ~100:1 (CFU:A549) was used for adherence and invasion to A549 cells **(A)** and ~5:1 (CFU:Airway Cell) for adherence to human airway cells **(B)**. 1000 CFU per well were used in neutrophil killing and opsonophagocytic killing assays **(C)**. Approximately 50 000 CFU per well were used in the whole blood killing assay **(D)**.
