## Supplementary Figure 2 for "A phasevarion controls multiple virulence traits, including expression of vaccine candidates, in *Streptococcus pneumoniae*"

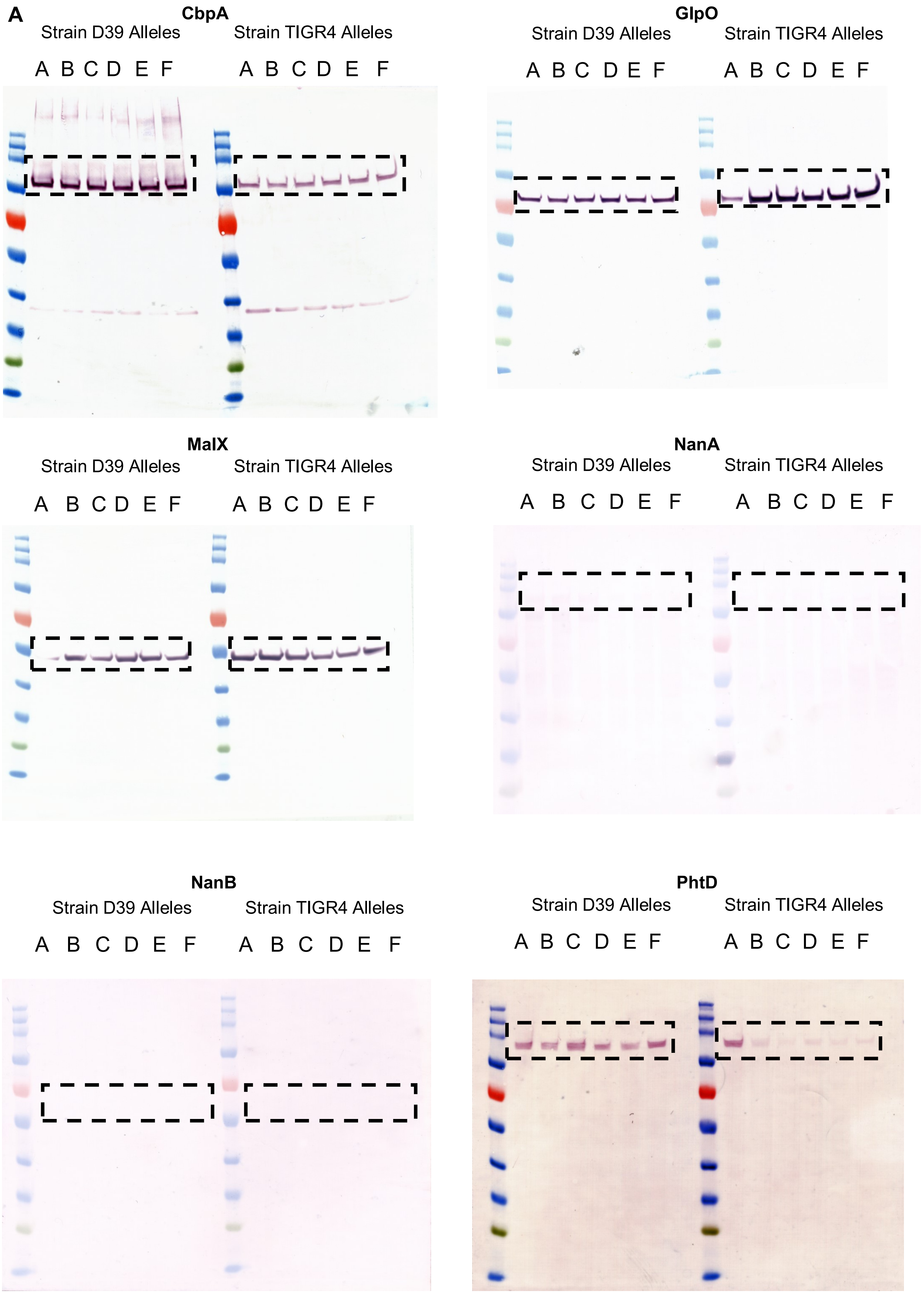
A
CbpA
Strain D39 Alleles
Strain TIGR4 Alleles
A B C D E F
A B C D E F
GlpO
Strain D39 Alleles
Strain TIGR4 Alleles
A B C D E F
A B C D E F
MalX
Strain D39 Alleles
Strain TIGR4 Alleles
A B C D E F
A B C D E F
NanA
Strain D39 Alleles
Strain TIGR4 Alleles
A B C D E F
A B C D E F
NanB
Strain D39 Alleles
Strain TIGR4 Alleles
A B C D E F
A B C D E F
PhtD
Strain D39 Alleles
Strain TIGR4 Alleles
A B C D E F
A B C D E F

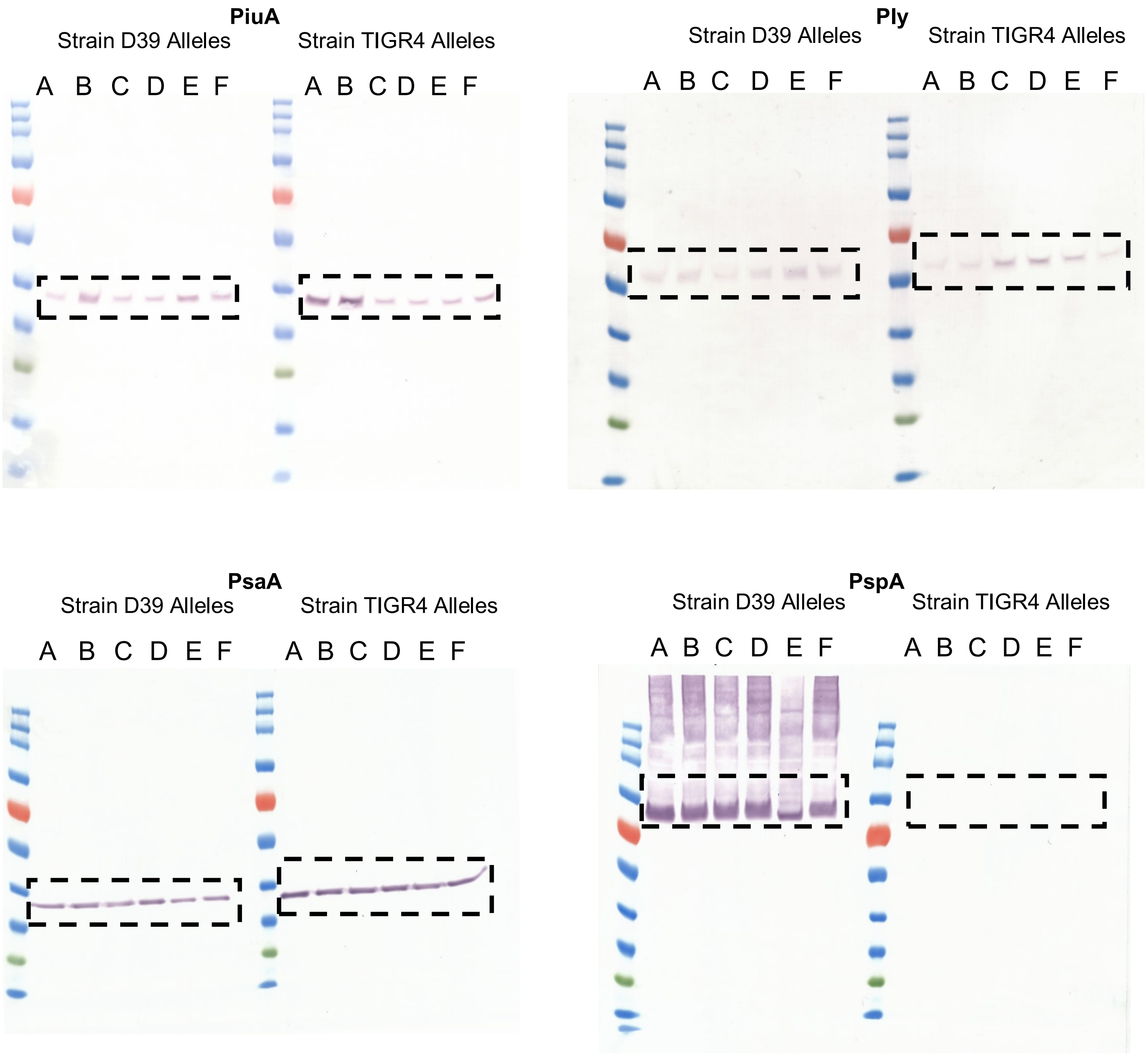


**Supplementary Figure 2A.** Full images of vaccine candidate Western Blots. Full scans of Western Blots using vaccine candidate anti-sera against *S. pneumoniae* D39 and TIRG4 locked SpnIII alleles. The outlined section of each Blot has been used in **Figure 3**.

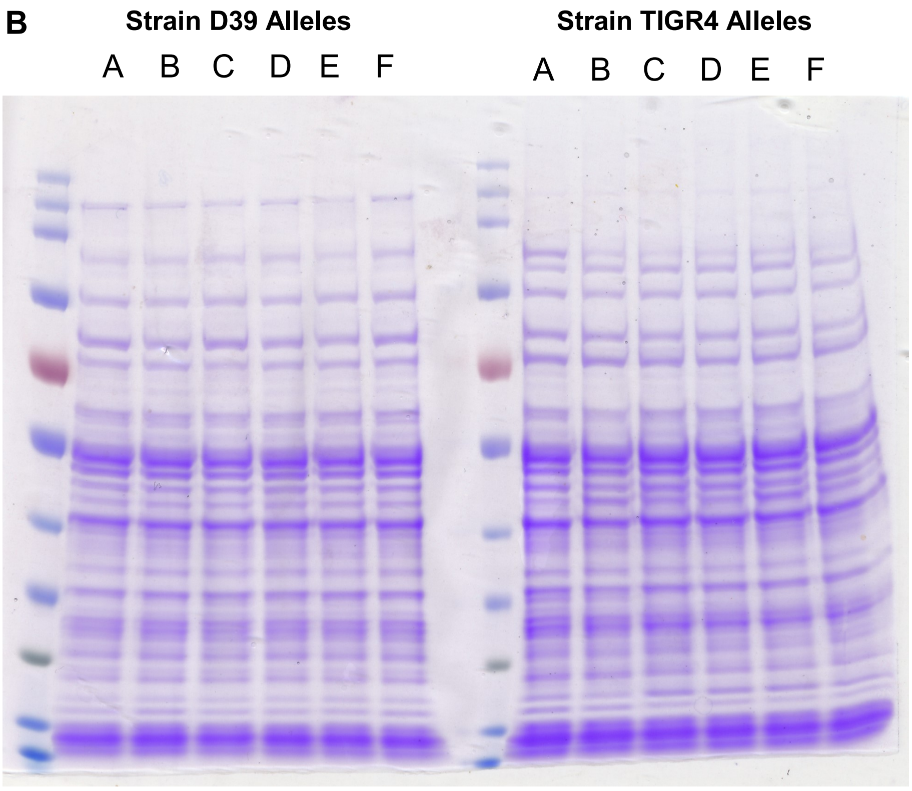


**Supplementary Figure 2B.** Balanced loading of locked allele lysates for Western Blots. D39 and TIGR4 locked allele cell lysate loads were standardised via Coomassie stain to ensure equal protein loading was used in vaccine candidate Western Blots used in **Figure 3** and **Supplementary Figure 2A**.

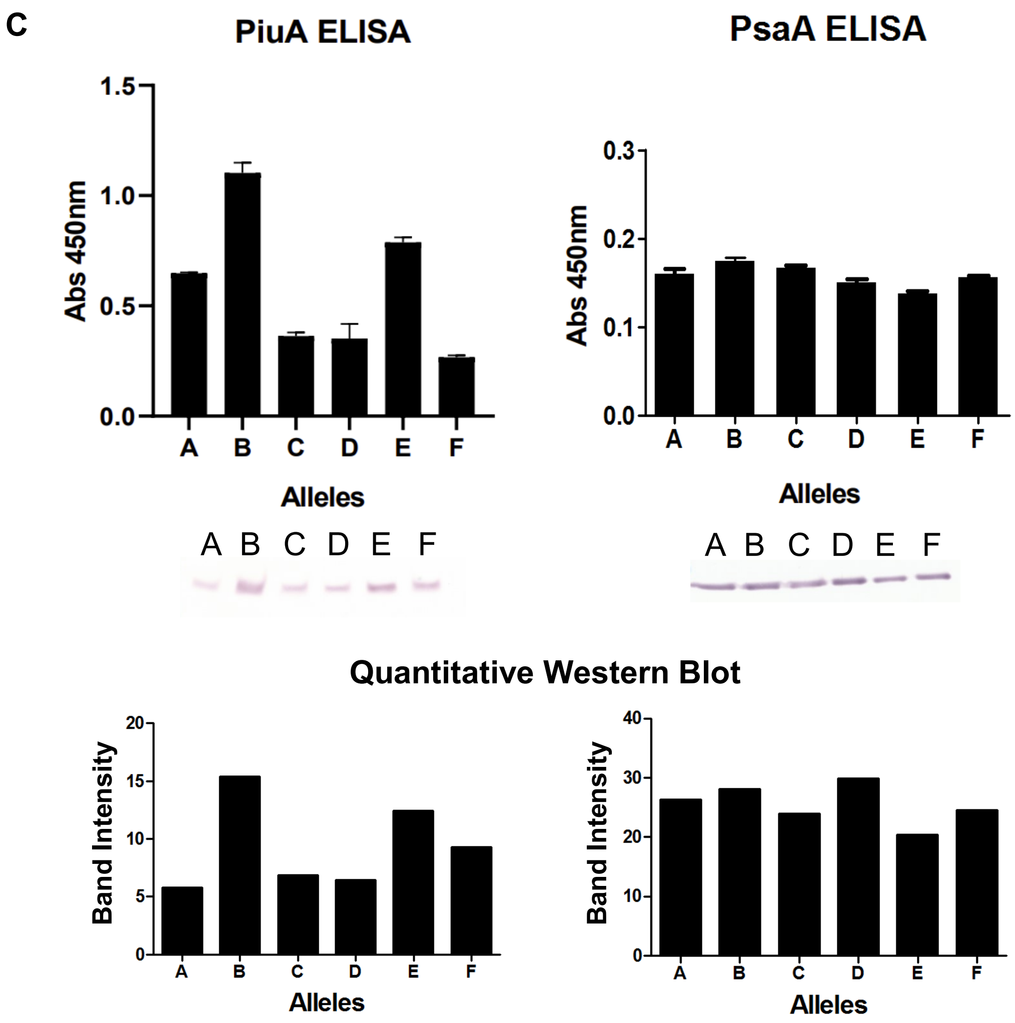


### Fold Difference from Most Intense Band

| PiuA Fold Difference | |
|---|---|
| B vs A | 2.7 |
| B vs C | 2.3 |
| B vs D | 2.4 |
| B vs E | 1.2 |
| B vs E | 1.7 |

| PsaA Fold Difference | |
|---|---|
| D vs A | 1.1 |
| D vs B | 1.1 |
| D vs C | 1.2 |
| D vs E | 1.5 |
| D vs F | 1.2 |

**Supplementary Figure 2C.** PiuA and PsaA ELISA with comparative Western Blot and quantified Western Blot values. Western Blot banding intensity was quantified in ImageJ. Fold differences calculated from the quantified Western Blot values are provided, using the most intense band as a baseline for comparison.
